# Whole body elongation drives coordinated vertebral shape evolution in Lake Malawi cichlid fishes

**DOI:** 10.64898/2026.05.09.723978

**Authors:** Callum V. Bucklow, Hannah Ugboma, Katharine E. Criswell, Mexford Mulumpwa, Roger Benson, Berta Verd

## Abstract

Understanding how anatomical structures evolve requires disentangling the roles of integration and modularity in shaping morphological variation. The vertebral column, a serially repeated and regionally differentiated structure, provides a powerful system for investigating these processes. Here, we examine how vertebral morphology evolves in relation to whole-body elongation across the adaptive radiation of Lake Malawi cichlid fishes. We tested for evolutionary integration between the precaudal and caudal domains, as well as assessed the contributions of vertebral count, centrum shape, and intervertebral spacing on body elongation. We find strong evolutionary integration between the shapes of precaudal and caudal vertebrae, with both vertebral shapes varying along similar axes. Despite this, precaudal and caudal vertebral counts evolve independently, indicating a decoupling between the specification of identity and the development of their respective shapes. Whole-body elongation is significantly associated with coordinated changes in vertebral and rib morphology, including proportional increases in centrum size, posterior displacement of neural and haemal spines, and increased rib curvature. In contrast, centrum elongation and intervertebral spacing do not contribute to body elongation across the radiation. These results demonstrate that body elongation in cichlids necessitates integrated, multivariate changes in axial morphology. Our findings highlight the importance of morphological integration in facilitating coordinated evolutionary responses in anatomical systems.

## INTRODUCTION

Morphological traits arise through the coordinated evolution of multiple interacting components, and the extent to which such traits are shaped by integration and modularity is a central question in evolutionary biology (Klingenberg, 2014; Wagner and Altenberg, 1996; Goswami et al., 2014). Integration, covariance amongst traits, can constrain evolutionary trajectories by coupling changes across structures, often reflecting shared developmental pathways and genetic interactions, but it can also channel evolution along coordinated pathways in response to selection (Pigliucci, 2003; Hallgrímsson et al., 2009), potentially allowing larger phenotypic changes than would be possible in the absence of integration (Goswami et al., 2014). In contrast, modularity enables components to evolve more independently, facilitating divergence among distinct organismal structures despite underlying developmental or functional relationships (Wagner and Altenberg, 1996; Klingenberg, 2014). Disentangling the developmental and evolutionary bases of integration and modularity is therefore critical for understanding phenotypic diversification (Goswami et al., 2014; Klingenberg, 2014; Orkney et al., 2021).

The vertebral column provides a powerful system for examining how anatomical structures evolve. As a defining feature of vertebrates, it plays a central role in locomotion, providing structural support, sites for muscle attachment, and protection for the spinal cord (Ford, 1937). As a serially repeated structure organised into distinct regions, it exhibits considerable variation in both form and function along the anterior-posterior axis, while retaining strong continuity among its constituent elements. This organisation reflects its developmental basis: vertebral count is determined by the developmental process of somitogenesis and the resulting number of somites that form, whereas vertebral identity is specified by anterior–posterior *Hox* gene expression in those somites (Morin-Kensicki et al., 2002; Woltering et al., 2009), with subsequent differentiation generating shape variation along the column (Apschner et al., 2011; Dietrich et al., 2020).

The vertebral column of teleost fishes is regionalised into precaudal and caudal regions (Figure 1A). Both vertebral types share a common structure comprising a neural spine and a centrum; the latter is formed by the fusion of anterior and posterior cone-shaped structures, which form a central canal that houses notochordal tissue. Precaudal vertebrae are characterised by basapophyses that support ribs and are associated with protection of the viscera and muscle attachment (Figure 1B), whereas caudal vertebrae possess haemal arches formed by haemal spines, which enclose a haemal canal housing the caudal artery and vein (Figure 1C). The neural and haemal spines of the caudal vertebrae articulate with the anal and caudal fins (Figure 1A), contributing to propulsion and manoeuvrability during swimming (Ford, 1937).

**Figure 1.**
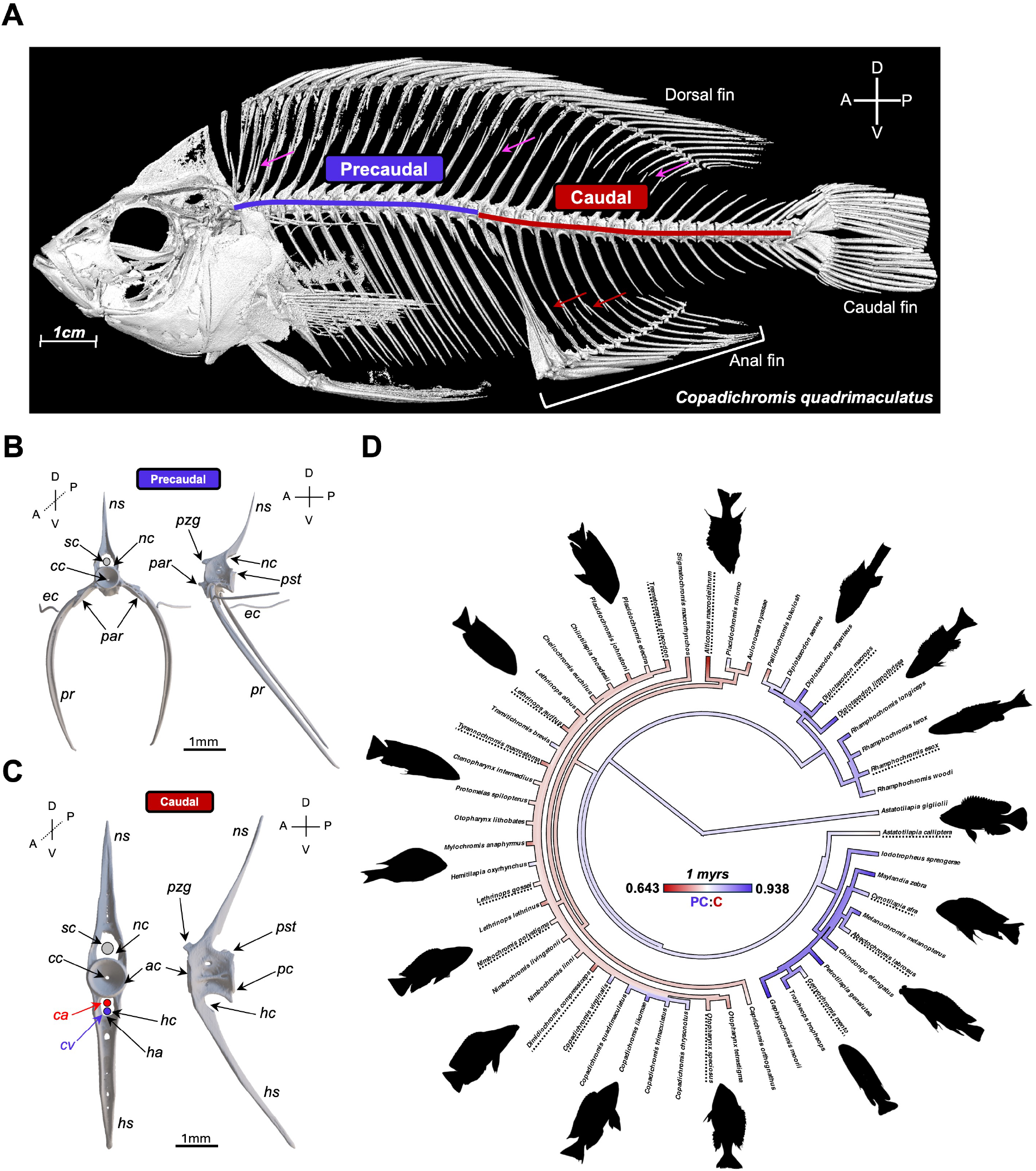
Anatomy of the axial skeleton and diversity in Lake Malawi cichlids. **(A)** A 3D rendering of a *µ*CT-scan of *Copadichromis quadrimaculatus*, for specimen details see Supplementary Data. The precaudal (purple) and caudal (red) vertebrae are indicated. Pink arrows indicate articulation of the neural spines with the dorsal fin rays (pterygiophores), which is common to precaudal and some caudal vertebrae. Red arrows indicate articulation of the haemal spines, associated with the caudal vertebrae, with the anal fin. Anatomical orientation axes are indicated (anterior–posterior, A–P; dorsal–ventral, D–V). Labelled models of precaudal **(B)** and caudal **(C)** vertebrae from *Maylandia zebra* are indicated, respectively. Abbreviations: *ac*, anterior cone; *ca*, caudal artery; *cc*, central canal; *cv*, caudal vein; *ec*, epicentral (dorsal rib); *ha*, haemal arch; *hc*, haemal canal; *hs*, haemal spine; *na*, neural arch; *nc*, neural canal; *ns*, neural spine; *par*, parapophyses; *pc*, posterior cone; *pr*, pleural rib; *pst*, postzygapophyses; *pzg*, prezygapophyses; *sc*, spinal cord. **(D)** Phylogeny showing branch-wise variation in precaudal–caudal vertebral ratio with inferred ancestral estimates, a lower value indicates greater relative caudal counts (more red). Silhouettes represent the body shapes of select species (dashed lines).

Vertebral count, proportions, and shape are highly variable across ray-finned fishes, ranging from the reduced axial skeleton of ocean sunfishes (*Mola*), with as few as 17 vertebrae and loss of the caudal fin in closely related species, to the extreme elongation and high vertebral counts of snipe eels (*Nemichthys*), which can exceed 700 vertebrae (Britz and Johnson, 2005; Deniz and Ağilkaya, 2018). In some lineages, such as syngnathids (seahorses, pipefishes, and seadragons), elements associated with precaudal vertebrae, including ribs and epicentrals, have been secondarily lost (Small et al., 2016; Schneider et al., 2023). This diversity highlights the evolutionary lability of the axial skeleton, but also raises the question of how changes in different vertebral components are coordinated.

Lake Malawi cichlids are a highly speciose and phenotypically diverse adaptive radiation of teleosts, comprising approximately 850 species (Turner et al., 2001; Genner and Turner, 2012). Despite their rapid diversification over ≈ 800 thousand years (Malinsky et al., 2018), genetic divergence among species is extremely low, with genome-wide sequence differences comparable to those observed within human populations (Svardal et al., 2021). This combination of high phenotypic diversity, low genetic variation and rapid divergence makes the radiation a powerful system for investigating morphological evolution (Santos et al., 2023). A major axis of this diversity is variation in body shape, particularly body elongation, which is closely associated with changes in vertebral column structure (Bucklow et al., 2025b). Increases in vertebral count are known to contribute to body elongation in teleosts (Ward and Brainerd, 2007; Mehta et al., 2010; Bucklow et al., 2025b), and substantial diversity in vertebral counts and identity, as well as body shape has evolved in Lake Malawi cichlids (Bucklow et al., 2024, 2025b; Oliver, 2024) (Figure 1D). However, far less is known about how vertebral shape evolves, or whether shape changes occur independently across regions of the axial skeleton. In particular, it remains unclear whether precaudal and caudal regions evolve as independent modules or as an integrated system and how the evolution of body elongation may be related to changes in vertebral shape.

Here, we quantify vertebral and pleural rib shape across Lake Malawi cichlids using three-dimensional geometric morphometrics to test how axial morphology evolves in relation to body elongation. Specifically, we (i) assess the degree of evolutionary integration between precaudal and caudal vertebrae, (ii) evaluate the contributions of vertebral counts, centrum shape, and intervertebral spacing to body elongation, and (iii) test whether elongation is associated with coordinated changes in multivariate vertebral shape. If vertebral regions evolve as independent modules, we expect weak covariation between precaudal and caudal morphology; alternatively, strong integration would indicate that body elongation drives coordinated vertebral shape changes across the axial skeleton.

## METHODS AND MATERIALS

### Geometric Morphometrics

#### Species Selection

We restricted the analysis to species present on the phylogeny of (McGee et al., 2020), resulting in vertebral shape data for 58 specimens, representing 53 species from across the radiation (Figure 1D). This subset spans all seven major ecomorphological groups described in Lake Malawi (Malinsky et al., 2018), capturing a wide range of axial skeletal variation within the radiation. Of the 58 specimens, 44 were previously published as part of a dataset of whole body *µ*CT-scans of Lake Malawi cichlids (Bucklow et al., 2024), eight additional specimens were downloaded from MorphoSource (Boyer et al., 2016) from the Yale Peabody Museum (YPM) collection whose *µ*CT-scans were collected by the oVert TCN (Blackburn et al., 2024), five belong to the Cambridge University Natural History Museum (CAMZM) collection and **a single *Maylandia zebra* from our live stocks that can be now found on Morphosource**.

#### Segmentation of µCT-scans

CT Image volumes were imported into Avizo for manual segmentation of vertebrae. Precaudal and caudal vertebral counts vary between African cichlids, including among Lake Malawi species (Oliver, 2024; Bucklow et al., 2025b) (Figure 1D). The first caudal vertebra was identified as the first haemal-arch bearing vertebrae, which articulates with the second anal fin spine (Supplementary Figure 1). The presence of ‘*transitional*’ vertebrae bearing both precaudal and caudal vertebral morphology (De Clercq et al., 2017) made defining the final precaudal and the first caudal vertebrae difficult (Supplementary Figure 1). Therefore, to ensure comparability between species, we segmented three precaudal and caudal vertebrae relative to the first haemal-arch bearing vertebrae, ignoring the vertebrae immediately anterior or posterior this articulation point (Supplementary Figure 1). Precaudal and caudal vertebral shapes were significantly different both within specimens and between species based on the shared, homologous landmarks (see Results).

#### Landmarking of Vertebral Models

The full landmark scheme is outlined in Supplementary Figure 2. We used a set of 15 landmarks (*n*_*L*_ = 15) common to both precaudal and caudal vertebrae, that captured centrum and neural spine shape variation. For the two-block PLS and whole body elongation analysis (see below), two additional landmarks were added to the caudal vertebrae to capture variation in haemal spine placement, which included the anterior fusion point of the haemal arch and the distal tip of the haemal spine. Although precaudal vertebrae and pleural ribs are anatomically distinct elements with different developmental origins (Woltering et al., 2018) and were analysed as separate structures, they were segmented and landmarked together within the same 3D models (see Supplementary Figure 2). To capture pleural rib curvature, two fixed landmarks were placed on the dorsal and ventral distal tips of the rib and ten semilandmarks equidistantly along the outer curve of the rightward pleural rib. Model files were imported into 3D-Slicer (v5.8.1) and landmarks were manually placed for all the models. Coordinate data for each landmark and specimen was exported as a .JSON file and compiled into a TPS (Thin Plate Spline) file in R for further analysis.

#### Linear Measures

We calculated both a mean centrum aspect ratio and intervertebral distance to assess their contribution to whole body elongation. For the anteroposterior centrum length, we took the mean of the natural logarithm of the Euclidean distances between corresponding landmarks on the anterior and posterior cones of the centra (landmarks 3-12, 5-14 and 4-13), respectively. The mean ln[length] was used as the centrum length for each respective vertebrae. Dorsoventral depths were calculated using the Euclidean distance between landmark pairs 3–4 and 12–13, that mark the depths of the anterior and posterior cones, respectively (see Supplementary Figure 2). The mean ln-transformed distance was taken and the aspect ratio was calculated by subtracting the mean depth from the mean length (ln[Centrum Length]-ln[Centrum Depth]). To estimate intervertebral distances (IVDs), Euclidean distances were calculated between adjacent cone landmarks, either posterior-to-anterior or anterior-to-posterior depending on vertebral orientation, using landmark pairs 3–12, 5–14, and 4–13. For each individual, we calculated the natural logarithm of each IVD and averaged these values within the precaudal and caudal regions separately. To obtain a single IVD value per species, the mean precaudal and mean caudal ln[IVD] values was taken. IVDs scaled positively with ln-transformed standard length (*R*^2^ = 0.424, *β*_1_ = 1.23, *λ* = 0.452, *p* < 0.0001, *d*.*f*. = 51). To account for this scaling effect, we calculated a specimen length-corrected IVD by subtracting ln[Specimen Length] from ln[IVD]. This removed the residual scaling effect (*R*^2^ = 0.026, *β*_1_ = 0.234, *λ* = 0.452, *p* = 0.249, *d*.*f*. = 51), permitting meaningful species comparisons. Whole body aspect ratios (ln[Length]-ln[Depth]) were calculated for each specimen in the dataset, as previously described (Bucklow et al., 2024) from volume renderings of the *µ*CT-scans.

### Generalised Procrustes Analysis

We used generalised Procrustes superimposition to remove variation in size, orientation and position among samples, using the *gpagen* function in the R package *geomorph* (v4.0.6) (Baken et al., 2021). Superimposition was performed separately on precaudal and caudal (*n*_*L*_ = 15) vertebrae and for the pleural ribs (*n*_*L*_ = 12). Alignments were iterated 100 times and were obtained by maximising Procrustes distance, with bending energy minimised for the pleural ribs to accommodate the lack of strict homology among landmarks (which were permitted to slide) along the curved rib. Since we had three bones for the precaudal, caudal and pleural ribs per specimen, the mean precaudal, caudal and pleural rib shape was calculated for each specimen from the aligned coordinates using the *mshape* function also in *geomorph* (v4.0.6) (Baken et al., 2021). To remove residual allometric scaling, we regressed the Procrustes-aligned mean coordinates against the mean log centroid size using the *procd*.*pgls* function in *geomorph* (v4.0.6) (Baken et al., 2021), which applies residual randomisation as implemented in the *RRPP* package (Collyer and Adams, 2018). Procrustes coordinates for all three bones were significantly correlated with log-centroid size (*p* < 0.01), therefore, we used the residuals from these multivariate regressions for all downstream analyses.

#### Principal Component Analysis

To identify the major axes of shape variation, we performed a principal component analysis (PCA) on the log-centroid size corrected shape residuals for the precaudal and caudal vertebrae (shared homologous landmarks, *n*_*L*_ = 15), as well as the pleural ribs (*n*_*L*_ = 12) using the *gm*.*prcomp* function also in the geomorph package (v4.0.6) (Baken et al., 2021). Because our goal was to explore overall shape variation rather than accounting for phylogenetic relatedness, we used standard PCA rather than a phylogenetically corrected PCA. To summarise the shape changes associated with PC1–PC5 (PC1–PC4 for the pleural ribs), we ranked coordinate loadings for each PC and extracted the five with the highest absolute values. These were interpreted alongside visualisation of shape variation along each PC axis by comparing the mean consensus shapes against the shapes ±2*σ* from the mean. A description of each PC axis, and the shape change it represents is indicated in Table 1.

**Table 1:** Summary of PC shape change axes. Shape change descriptions given represent the change from low to high PC value. *PC change is the same.

| PCA Axis | Bone | Variance Explained (%) | Shape Change |
| --- | --- | --- | --- |
| 1 | Precaudal | 47.47 | *Ventral and posterior shift in the position of the neural spine tip. Proportionally larger centra. Neural spines proportionally shorter.\** |
|  | Caudal | 54.32 | *Ventral and posterior shift in the position of the neural spine tip. Proportionally larger centra. Neural spines proportionally shorter.\** |
|  | Pleural Rib | 54.71 | Increase in rib curvature, ventral tip of the pleural rib is shifted posteriorly and dorsally. |
| 2 | Precaudal | 16.16 | *Anterior shift in neural spine tip but with corresponding posterior and ventral shift in tip of the neural arch fusion point. Neural spine appears 'straighter' and neural arch is 'swept back'.<sup>+</sup> Lateral and posterior shift in the posterior insertion of the neural spine into the centrum representing a widened point of fusion between neural spine and centrum.* |
|  | Caudal | 14.75 | *Anterior shift in neural spine tip but with corresponding posterior and ventral shift in tip of the neural arch fusion point. Neural spine appears 'straighter' and neural arch is 'swept back'.<sup>+</sup> Ventral shift in posterior cone relative to the anterior cone.* |
|  | Pleural Rib | 17.88 | Rib curve is shifted ventrally |
| 3 | Precaudal | 10.73 | Dorsal shift of anterior cone and ventral shift in relative position of the posterior cone (relative to anterior cone), 'straightening' of the relative cone positions. Dorsal shift in the position of the anterior fusion point of neural spine and centrum. |
|  | Caudal | 8.34 | Dorsal shift in the insertion of the neural spine into the centrum (i.e. appears to be higher up around the centrum), ventral shift in the posterior insertion point of the neural spine with centrum (i.e. a shorter fused section). |
|  | Pleural Rib | 11.51 | Lateral (outward) shift in the pleural rib curve but ventral portion of rib is relatively straight. |
|  | Precaudal | 5.93 | Anterior shift in the posterior contact between centrum and neural arch. |

Table 1: **Summary of PC shape change axes.** Shape change descriptions given represent the change from low to high PC value. \*PC change is the same. (Continued)
|  |  |  |  |
| --- | --- | --- | --- |
|  | Caudal | 4.51 | Anterior shift in posterior insertion/fusion point of neural spine with centrum, representing a thinning of the neural spine fusion/insertion. |
|  | Pleural Rib | 5.24 | Lateral and posterior shift in the dorsal tip and dorsal portion of pleural rib curve. |
| 5 | Precaudal | 4.94 | Posterior and ventral shift in ventral midpoint of centrum, representing a shallower depression in the ventral portion of the centrum. |
|  | Caudal | 3.48 | Posterior shift in the posterior cone, perhaps reflecting elongation of centra and a dorsal shift in the depression formed on the ventral side of the centrum, representing a thinning of the middle proportion of the centrum. |
|  | Pleural Rib | N/A | N/A |

## Phylogenetic Comparative Methods

### Phylogenetic Linear Models

To evaluate the relationships among evolutionary changes in centrum aspect ratio, intervertebral distance (IVD), total vertebral counts, and whole-body aspect ratio, we used phylogenetic generalised least squares (PGLS; (Grafen, 1989)) implemented in the *pgls* function of the R package *caper* (v1.0.1), which reports phylogenetically informed *R*^2^ values (Orme et al., 2018). To account for phylogenetic signal in trait covariance, we estimated Pagel’s *λ* branch transformation parameter by maximum likelihood. Pagel’s *λ* rescales the expected covariance among species under a Brownian motion model of evolution: *λ* = 1 corresponds to the covariance structure expected under Brownian motion along the phylogeny, whereas *λ* = 0 implies statistical independence among species, such that trait relationships are not structured by shared ancestry (Pagel, 1997, 1999).

To test for the co-evolution of precaudal and caudal shape variation and body elongation, we used multivariate phylogenetic regression (Adams, 2014). Since our two-block partial least squares analysis indicated strong integration between precaudal (*n*_*L*_ = 15) and pleural rib (*n*_*L*_ = 12) shape, as well as between precaudal (*n*_*L*_ = 27, including pleural ribs) and caudal vertebral shape (*n*_*L*_ = 17, including haemal spines), we used the residuals of Procrustes coordinates regressed against log-transformed centroid size (as described above). Residuals were regressed against the specimen ln[Length]-ln[Depth], using the *procD*.*pgls* function in *geomorph* (v4.0.6) (Baken et al., 2021). Statistical significance was assessed using 10,000 permutations with residual randomisation as implemented in the *RRPP* package (Collyer and Adams, 2018). To assess the robustness of these regressions, we excluded species with body aspect ratio *z*-scores exceeding ±1.5 (i.e., all *Rhamphochromis, Diplotaxodon limnothrissa*, and *Lethrinops gossei*) and repeated the analyses on this reduced dataset.

### Two-block Partial Least Squares

To assess whether shape changes were integrated, we used a phylogenetic implementation of two-block partial least squares (Rohlf and Corti, 2000), a method that evaluates whether shape covariation between two structures is greater than expected under a Brownian motion model of evolution (p2BPLS; (Adams and Felice, 2014)). If a significant amount of covariation is detected between the two bones, it suggests that their shape changes are evolutionarily coordinated. Specifically, we tested the co-evolution of precaudal vertebrae (*n*_*L*_ = 15) and pleural rib (*n*_*L*_ = 12) shape, and separately between precaudal (*n*_*L*_ = 27, with pleural ribs) and caudal vertebrae (*n*_*L*_ = 17, with haemal spine landmarks) using the log-centroid size-corrected residuals (see above) using the *phylo*.*integration* function in *geomorph* (v4.0.6) (Baken et al., 2021). In the second analysis, pleural ribs were included with precaudal vertebrae because we found that rib and precaudal shape were moderately integrated (see Results). To determine whether whole-body elongation predicts coordinated shape change in precaudal and caudal vertebrae, precaudal and caudal PLS1 scores were first standardised (*z*-transformed to mean = 0, SD = 1) to account for differences in scale between blocks, and then averaged to generate a single metric of coordinated variation per species. We then fit a PGLS model with this composite PLS1 score as the dependent variable and body aspect ratio (ln[Length] – ln[Depth]) as the predictor. A significant relationship indicates that body elongation is associated with systematic variation along the primary axis of coordinated shape change between precaudal and caudal vertebrae.

## Data and Code Availability

All code used in the analysis has been deposited on GitHub and can be accessed here. 53 of the 58 of the *µ*CT-scans used in the study are already available on Morphospace (see Supplementary Data). Please contact authors directly for the remaining six. Precaudal and caudal count data is from a previous study (Bucklow et al., 2026a). Vertebral models, coordinate data, lateral images of whole bodies, specimen and species data has been deposited on Zenodo (Bucklow et al., 2026b).

## RESULTS

### Precaudal and caudal vertebrae have distinct but overlapping patterns of shape variation

Mean precaudal and caudal vertebral shape differed both within specimens (Procrustes ANOVA: *R*^2^ = 0.231, *F*(1, 51) = 106.87, *Z* = 4.24, *p* = 0.0001) and between species (*R*^2^ = 0.659, *F*(51, 51) = 5.99, *Z* = 13.01, *p* = 0.0001), indicating that precaudal and caudal vertebrae represent distinct morphologies and that vertebral shape varies significantly between species. Precaudal vertebral shape differs from that of caudal vertebrae by exhibiting greater neural arch displacement, a more posterior articulation between the neural spine and the centrum and a ventrally shifted anterior cone (Figure 2A). In addition, precaudal centra are also consistently more elongate than caudal centra across species (phylogenetic linear-mixed model; *β*_caudal_ = −0.066, *SE* = 0.0078, *p* < 0.0001), corresponding to an an approximately 6.4% lower ratio in caudal centra relative to precaudal centra (Figure 2B).

**Figure 2.**
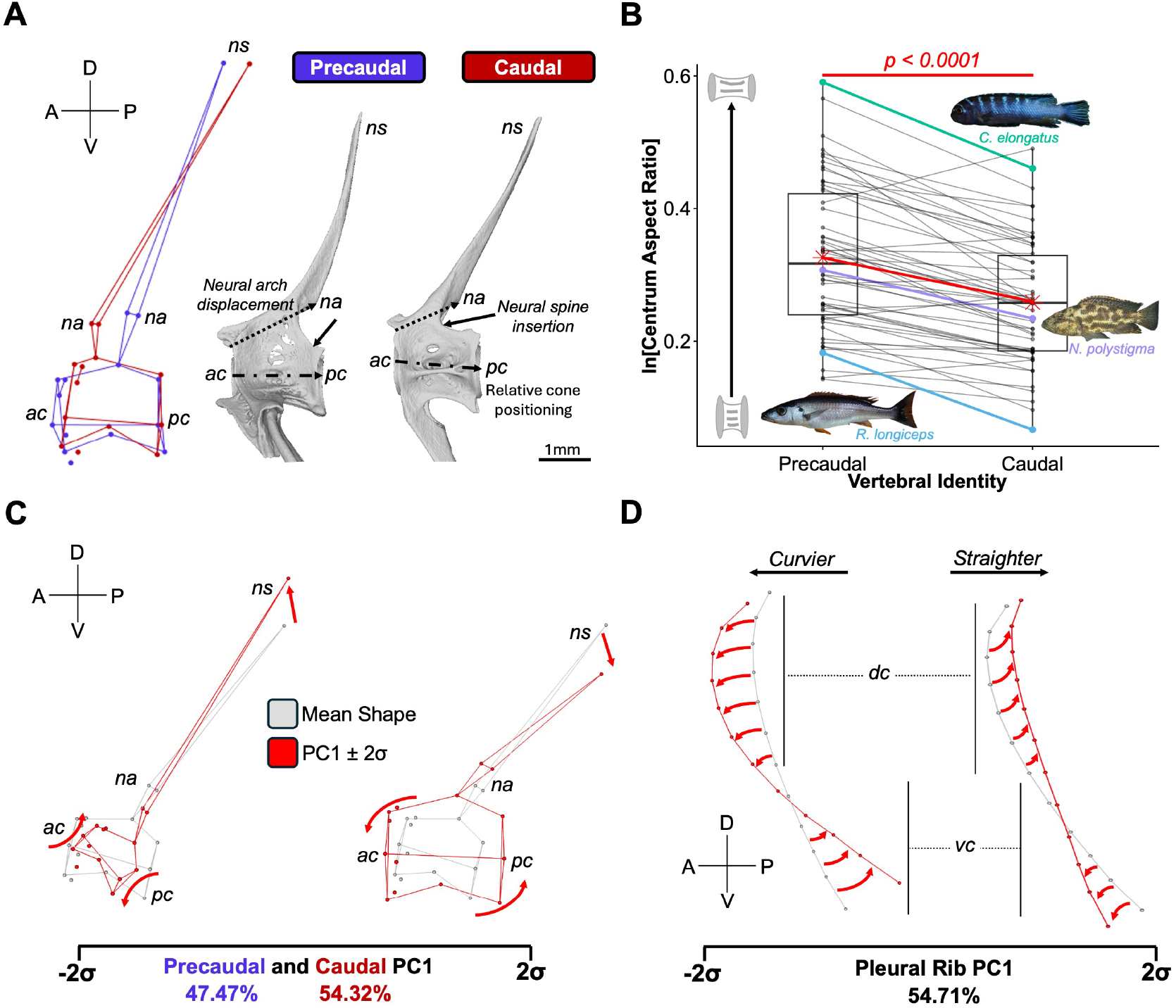
Precaudal and caudal vertebrae show distinct but overlapping patterns of shape variation. **(A)** Shape differences between precaudal and caudal vertebrae based on homologous landmarks capturing variation in the centra and neural spine. Mean shapes are overlaid to highlight differences. Anatomical orientation axes are indicated (anterior–posterior, A–P; dorsal–ventral, D–V). Representative precaudal and caudal vertebrae (*Maylandia zebra*) illustrate major shape features. Scale bar: 1 mm. **(B)** Boxplot of species mean ln-transformed centrum aspect ratios for precaudal and caudal vertebrae. Lines connect mean precaudal and caudal centrum aspect ratios (as ln[Centrum Length]-ln[Centrum Depth]) for individual species. Precaudal and caudal means are plotted as red stars. The overall mean difference is significant (p<0.0001), as indicated by a phylogenetic linear-mixed model (see Results). **(C)** PC1 is shared between precaudal and caudal vertebrae (see Table 1). The consensus precaudal shape (grey) is overlaid with shapes at ± 2*σ* along PC1. Red arrows indicate major deviations from the consensus. Percentage variation explained by PC1 is shown on the axis. Note that the figure only shows precaudal shape differences. **(D)** As in (C), but for pleural ribs. Abbreviations: *ac*, anterior cone; *dc*, dorsal curve; *na*, neural arch; *ns*, neural spine; *pc*, posterior cone; *vc*, ventral curve; *C., Chindongo*; *N., Nimbochromis*; *R., Rhamphochromis*. See supplementary information for image credits.

The first five principal component axes (PC1-5) explained a cumulative 84.24% (precaudal) and 85.40% (caudal) vertebral shape variation, respectively (see Table 1). PC1 accounted for considerable portions of shape variation for precaudal (PC1: 47.47%) and caudal (PC1: 54.32%) vertebrae, primarily reflecting proportional scaling of the vertebral centra, the relative length and positioning of the neural spine, and the positioning of the anterior and posterior cones in both (Figure 2C). For the pleural ribs, PCs 1–4 collectively represented 89% of the shape variation. PC1 accounted for 54.71% of the variation, primarily describing pleural rib curvature and the positioning of the ventral distal tip (see Table 1, Figure 2D).

### Vertebral shape evolution is strongly integrated, despite decoupled count evolution

Covariation between precaudal vertebral shape and the pleural rib shape was high. We found positive correlations between PLS1-PLS4 for precaudal and pleural rib shape covariation which jointly accounted for 97% of the total covariance between the two bones (r-PLS=0.629-0.518, all *p* < 0.0001). PLS1 accounted for 71% of the total covariance (r-PLS = 0.585, *Z* = 2.61, *p* < 0.0001, Figure 3A), which is primarily associated with dorsoventral shifts in the anterior and posterior cones, anteroposterior shifts in neural spine positioning and coordinated changes in the pleural rib dorsal and ventral arcs (Figure 3B). Therefore, rib curvature has been tightly integrated with evolutionary variation in precaudal vertebral shape.

**Figure 3.**
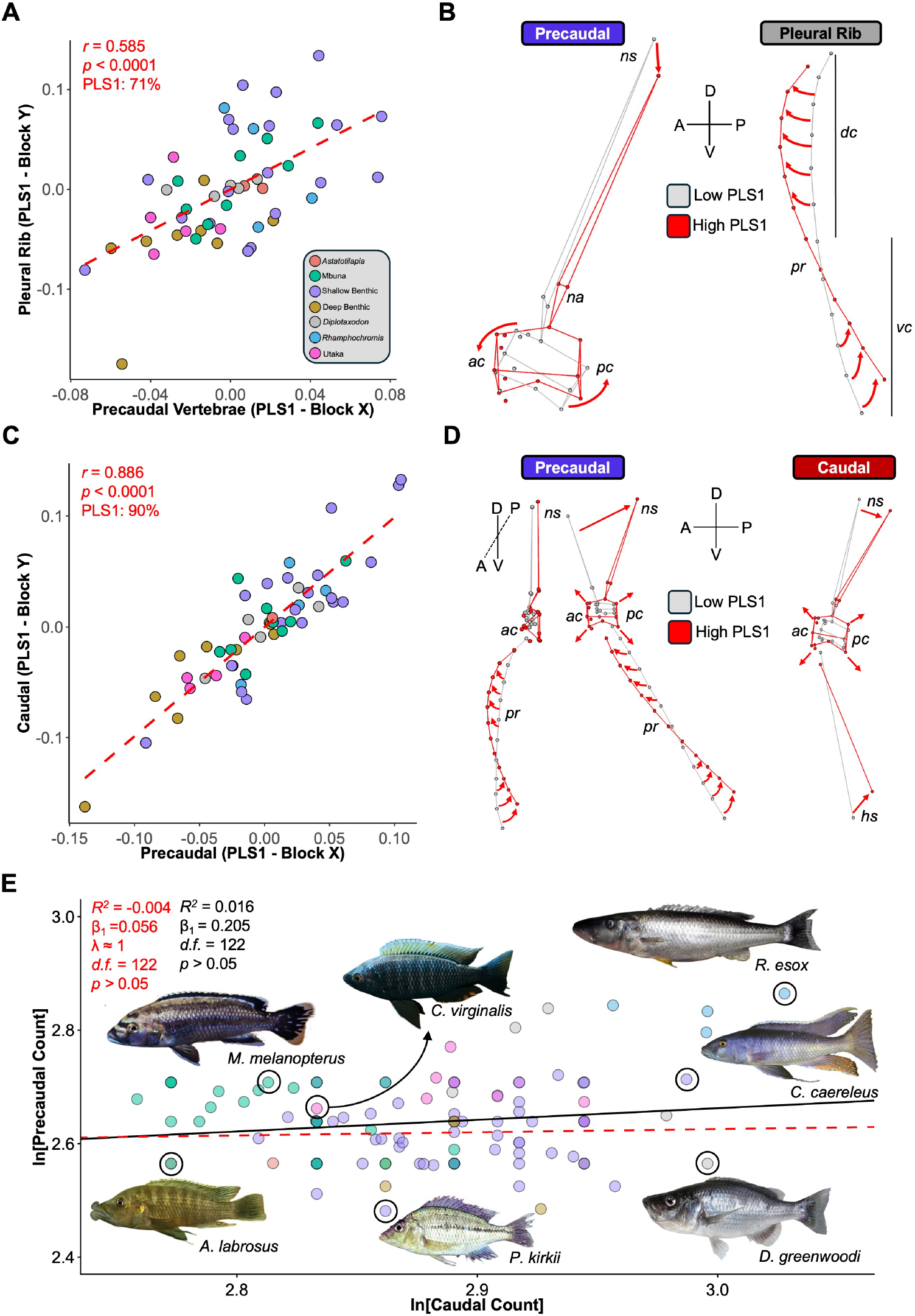
Vertebral shape evolution is strongly integrated despite decoupled count evolution. **(A)** Scatterplot of precaudal vertebrae and pleural rib PLS1 scores (blocks X and Y). Points represent species, coloured by ecomorphological group (see legend). **(B)** Shape change along PLS1 for precaudal vertebrae and pleural ribs. Low PLS1 is shown in grey, high PLS1 in red; arrows indicate change from low to high. Axes denote anatomical orientation (anterior–posterior, A–P; dorsal–ventral, D–V). **(C)** As in (A), for precaudal vs. caudal vertebrae. **(D)** As in (B), for precaudal (including pleural ribs) and caudal vertebrae (including haemal spines); an additional precaudal view highlights rib shape change. **(E)** Scatterplot of ln-transformed precaudal and caudal counts with ordinary least squares (black solid) and phylogenetic generalised least squares (red dashed) fits. Note that multiple points are overlapping. Abbreviations: *ac*, anterior cone; *dc*, dorsal curve; *na*, neural arch; *ns*, neural spine; *pc*, posterior cone; *vc*, ventral curve. Genera abbreviations: *A., Abactochromis*; *C., Copadichromis* (virginalis) or *Champsochromis*; *D., Diplotaxodon*; *M., Melanochromis*; *P., Protomelas*; *R., Rhamphochromis*. See supplementary information for image credits.

Precaudal vertebral shape (including the pleural ribs) is also strongly integrated with caudal vertebral shape. We found a strong positive correlation between block scores for PLS1 (r-PLS = 0.886, *Z* = 5.24, *p* < 0.0001, Figure 3C), which itself represented 90.18% of the precaudal-caudal shape covariation. Along PLS1, covarying shape changes closely resemble those of PC1, primarily representing proportional scaling of precaudal and caudal centra, anteroposterior positioning of the neural spine (and now haemal spine in caudal vertebrae), with proportional increases in centrum size associated with increases in pleural rib curvature (Figure 3D). Therefore, the evolution of vertebral shapes was highly integrated and coordinated during Lake Malawi cichlid diversification. Despite this strong shape integration, we did not detect a significant relationship between precaudal and caudal vertebral counts across the radiation (*R*^2^ = −0.004, *β*_1_ = 0.056, *λ* ≈ 1, *p* = 0.468, *d*.*f*. = 122; Figure 3E), consistent with previous findings in the subfamily (Bucklow et al., 2025b). Therefore, although precaudal and caudal vertebral shapes are highly integrated, their numbers appear to evolve independently, suggesting counts, but not their respective shapes, are evolutionarily decoupled.

### Whole body elongation is associated with coordinated changes in vertebral and rib morphology

We next asked whether variation in overall body form could account for the major axis of coordinated vertebral shape change. A significant proportion of variation in precaudal (*R*^2^ = 0.077, *F*(1, 50) = 4.21, *Z* = 2.69, *p* = 0.003; Figure 4A) and caudal (*R*^2^ = 0.087, *F*(1, 50) = 4.78, *Z* = 2.52, *p* = 0.003; Figure 4B) shape was explained by body elongation. This relationship remained significant after accounting for standard length, which showed no significant association with vertebral shape (both *p* > 0.05), confirming that the effect of body elongation is independent of overall body length. Therefore, body elongation is associated with proportionally larger vertebral centra, posteriorly and ventrally shifted neural and haemal spines, and increased anteroposterior and lateral curvature of the pleural ribs (Figure 4A, B), closely mirroring the shape changes captured by PC1 (Figure 2C) and PLS1 (Figure 3D). These multivariate shape changes are robust to the removal of taxa with extreme phenotypes, such as deep-bodied *Lethrinops* or elongate *Rhamphochromis*. Although *R*^2^ values and effect sizes were reduced (precaudal *R*^2^ = 0.057, *Z* = 1.98; caudal *R*^2^ = 0.069, *Z* = 2.11), both relationships remained significant (*p* < 0.05). Notably, standard length showed a modest but significant association with precaudal shape in this reduced dataset (*R*^2^ = 0.049, *F*(1, 45) = 2.35, *Z* = 1.74, *p* = 0.042), suggesting a secondary effect of body size that becomes detectable when the influence of extreme aspect ratios are removed. Nonetheless, extreme phenotypes strengthen but do not generate the observed relationship between elongation and vertebral shape. Consistent with the alignment between elongation-associated shape change and these major axes of (co-)variation, whole-body elongation explained a modest but significant proportion of covariation between precaudal (including pleural ribs) and caudal vertebral shape (including haemal spines), as captured by a standardised composite of their PLS1 scores (*R*^2^ = 0.09, *β*_1_ = 1.91, *λ* ≈ 0, *p* = 0.02, *d*.*f*. = 50; Figure 4C). Therefore, whole body elongation does not merely predict shape variation within precaudal and caudal vertebrae independently; it also explains a significant component of their shared covariation, implicating elongation as a key driver of integrated axial skeletal evolution during the diversification of Lake Malawi cichlids.

**Figure 4.**
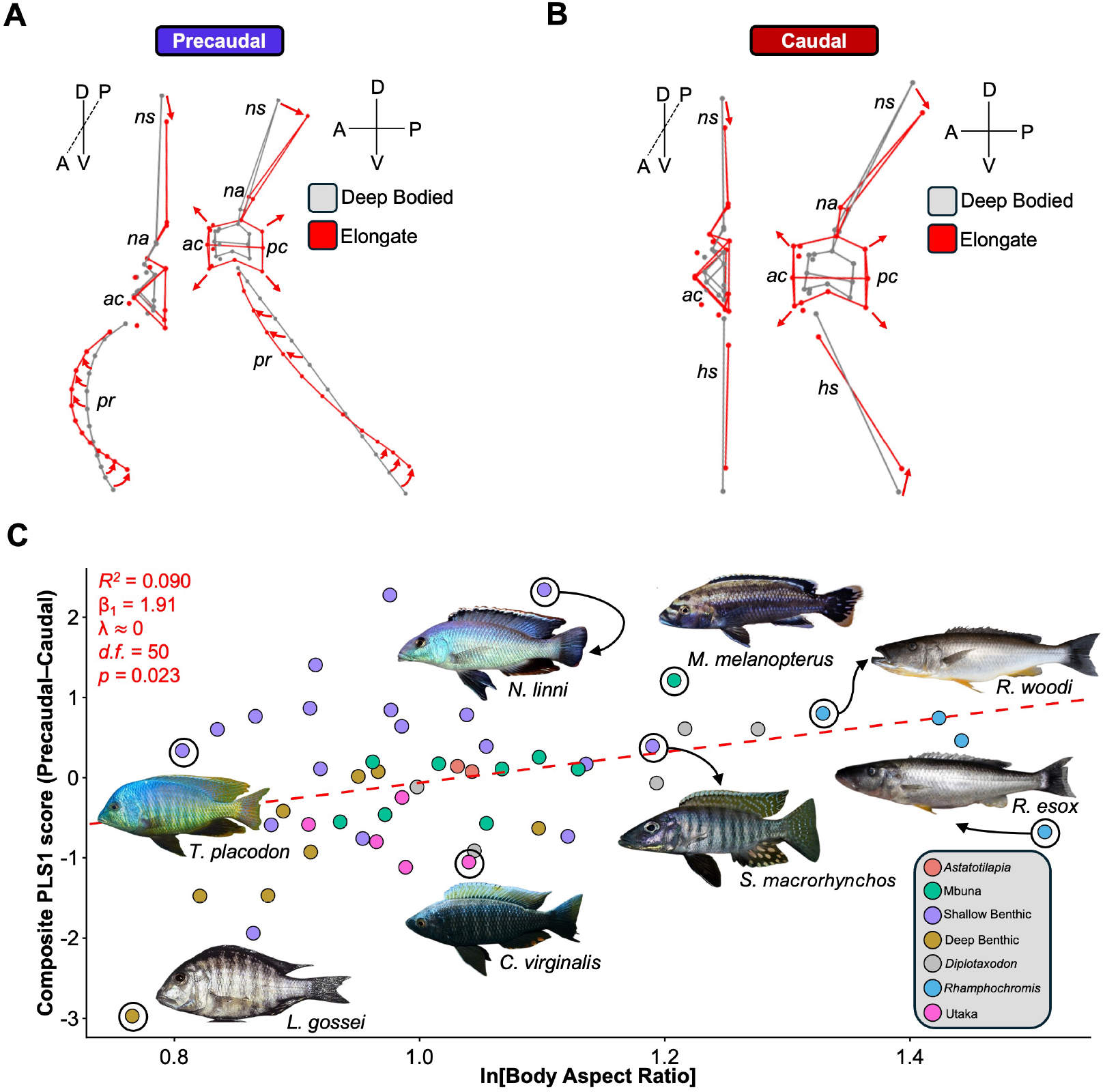
Whole body elongation is associated with coordinated changes in vertebral and rib morphology. **(A)** Precaudal shape change along the elongation axis. The grey shape represents the predicted morphology at minimum body elongation, contrasted with the red shape at maximum elongation. Red arrows indicate shape changes relative to the maximally elongated condition. **(B)** As in (A), but for caudal vertebrae. **(B)** as in (A) but for caudal vertebrae. Anatomical orientation axes are indicated (anterior–posterior, A–P; dorsal–ventral, D–V). **(C)** A scatterplot of standardised (z-score) block scores from the first PLS axis, averaged to obtain a single composite score per species for precaudal-caudal covariation against whole body elongation (as ln[Length]-ln[Depth]). The PGLS fit is indicated (dashed red line). OLS fit has been omitted as *λ* ≈ 0. Representative species are indicated. Abbreviations: *ac*, anterior cone; *dc*, dorsal curve; *na*, neural arch; *ns*, neural spine; *pc*, posterior cone; *vc*, ventral curve. Genera abbreviations: *C., Copadichromis* (virginalis) or *Champsochromis*; *L., Lethrinops*; *M., Melanochromis*; *N., Nimbochromis*; *R., Rhamphochromis*; *S., Stigmatochromis*; *T., Trematocranus*. See supplementary information for image credits.

### Centrum elongation and intervertebral spacing do not drive body elongation

To determine whether body elongation reflects simple geometric scaling or coordinated vertebral shape change, we tested whether centrum elongation and intervertebral spacing contribute to variation in body elongation beyond total vertebral counts. We found a significant positive correlation between total vertebral count and body elongation in Lake Malawi cichlids (Figure 5). The estimated slope for the Lake Malawi radiation (*β*_1_ = 1.77 ± 0.27 SE) is similar to that previously reported for the subfamily ((Bucklow et al., 2025b); Δ*β*_1_ = 0.31 ± 0.30 SE, *p* = 0.30), suggesting the effect of vertebral addition on body elongation within the radiation is consistent with the rest of the subfamily. However, we found no evidence that centrum aspect ratios or intervertebral distances have contributed to body elongation in the Lake Malawi radiation (Figure 5). An additive model including ln[Total Count], centrum aspect ratios and scaled IVDs (*R*^2^ = 0.333, *λ* = 0.648, *d*.*f*. = 49) was significant (*p* = 0.0002). However, only ln[Total Count] had a significant, positive slope (*β*_1_ = 1.47, p = 0.0004), with both centrum aspect ratios and intervertebral distances reporting non-significant negative slopes. Moreover, we found the additive model to be a considerably worse fit (ΔAICc = 4) than a model with ln[Total Count] alone (*R*^2^ = 0.339, *λ* = 0.603, *β*_1_ = 1.64, *p* < 0.0001, *d*.*f*. = 50). These results indicate that the relationship between whole body elongation and vertebral morphology is not mediated by simple changes in centrum shape or spacing, but instead reflects broader, coordinated changes in multivariate vertebral shape and in vertebral count evolution.

**Figure 5.**
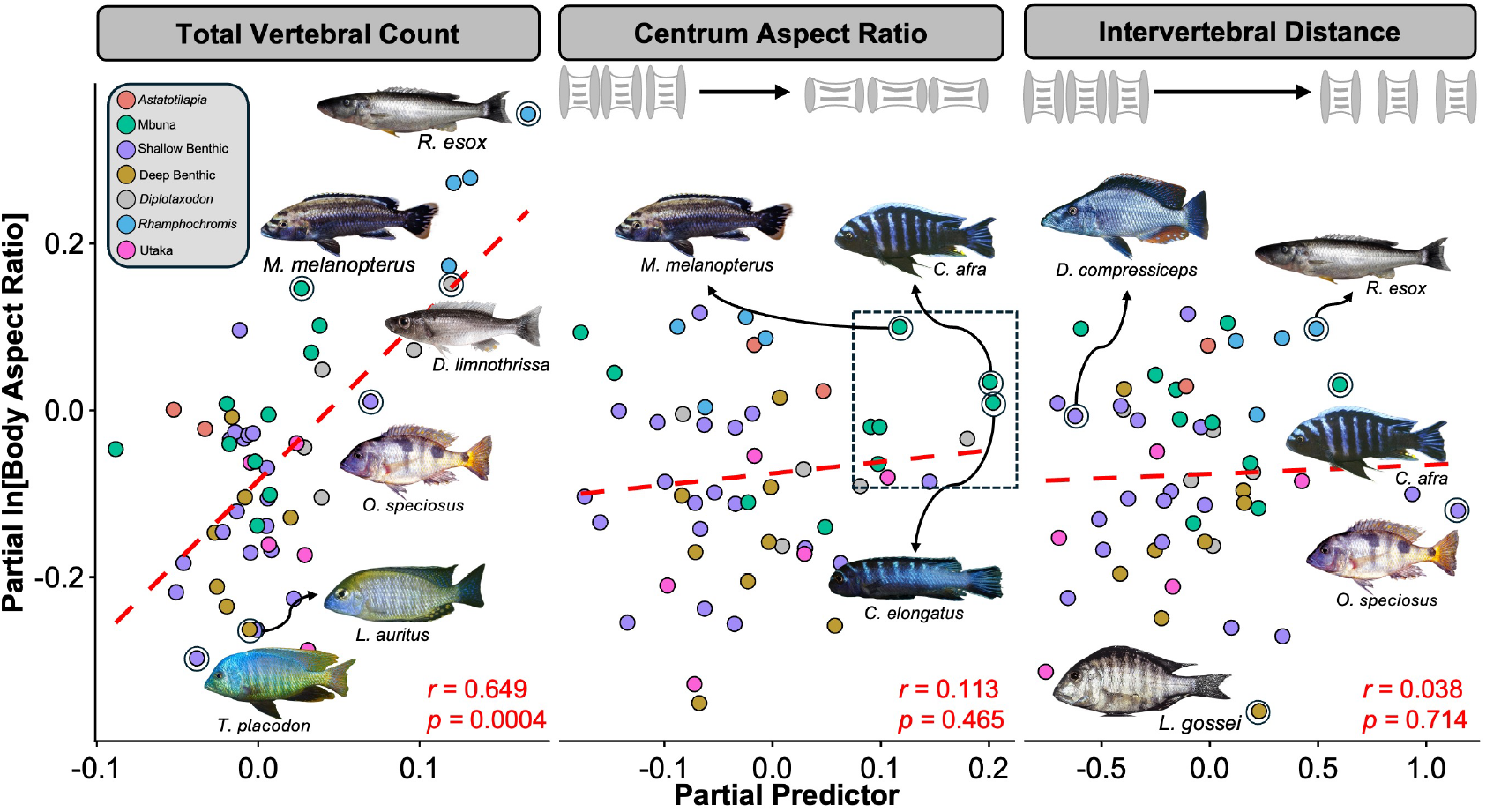
Centrum elongation and intervertebral spacing do not drive body elongation. Phylogenetic partial regression residuals for ln[Total Vertebral Count] (left), centrum aspect ratio (middle), and scaled intervertebral distances (right, relative to specimen standard length) against whole body elongation (as ln[Length]-ln[Depth]). Residuals represent the contribution of each variable within the full additive model. Correlation coefficients (*r*) indicate the strength of association between residuals; p-values report the significance of slopes from the full additive model. Note that slopes for the partial regressions of centrum aspect ratio and IVD are positive in the partial regression but negative in the full additive model (see Results). This occurs because shared variance with other predictors, accounted for in the full additive model, is absent in the partial regressions. Points are coloured by ecomorphological category (see legend), with images shown for selected species. Genus abbreviations: *C*. (*Chindongo*); *D*. (*Dimidiochromis*); *L*. (*Lethrinops*); *M*. (*Melanochromis*); *S*. (*Stigmatochromis*); *T*. (*Tropheops*); *R*. (*Rhamphochromis*).

## DISCUSSION

Whole-body elongation in Lake Malawi cichlids is associated with coordinated, multivariate changes across the axial skeleton. Precaudal, pleural ribs and caudal vertebrae, are strongly evolutionarily integrated and vary along shared shape change axes, with elongation predicting proportional increases in centrum size, posterior displacement of neural and haemal spines, and increased pleural rib curvature (Figure 4). This coordination likely reflects a combination of geometric and functional constraints imposed by changes in body shape. As bodies become more dorsoventrally compressed (elongate), neural and haemal spines likely shift to accommodate reduced vertical space, whereas deeper-bodied forms require proportionally longer and straighter neural spines and pleural ribs to span a wider body cavity. Along the elongation axis, changes in centrum organisation, particularly the relative positioning of the anterior and posterior cones, may also reflect adjustments in overall spinal curvature, with more elongate species exhibiting a relatively straighter vertebral column. Despite this strong integration of the shapes of the precaudal and caudal vertebral shapes, vertebral counts are the primary predictor of elongation, with no additional contribution from centrum elongation or intervertebral spacing. Therefore, whilst increasing vertebral counts are necessary to drive body elongation, vertebral morphology is constrained to evolve along shared, integrated trajectories. More broadly, these findings demonstrate that adaptive diversification in Lake Malawi cichlids is not an unconstrained exploration of morphospace, but has been channelled along, perhaps developmentally and functionally, integrated axes of variation.

Precaudal and caudal vertebral counts evolve independently, consistent with patterns across Pseudocrenilabrinae (Bucklow et al., 2025a), even though vertebral shapes do not. Homeotic shifts in the precaudal–caudal boundary are likely mediated by changes in *hox* gene regulation during somitogenesis, but identifying the key genes in teleosts is complicated by multiple *Hox* clusters (Crow et al., 2006; Hoegg et al., 2007) which encode paralogous genes with overlapping expression and functional redundancy (Adachi et al., 2024). Moreover, subfunctionalisation (Scemama et al., 2006; Le Pabic et al., 2007), lack of developmental conservation (Sordino et al., 1995; Adachi et al., 2024), and potential epigenetic regulation (Xue et al., 2022), suggests that the specific *hox* genes involved likely vary among teleosts, highlighting the need for comparative studies of *hox* expression. In addition, the precaudal and caudal domains may comprise multiple, independent subdomains (Jawad et al., 2018; Woltering et al., 2018), likely defined by distinct *hox* expression boundaries (Criswell et al., 2021), allowing fine-scale changes in vertebral counts without coordinated shifts across the entire axial skeleton. Nonetheless, although precaudal and caudal counts can evolve independently (Figure 3E) (Ward and Mehta, 2010; Bucklow et al., 2025a), the strong integration of precaudal and caudal shape suggests a separate mechanism acting downstream of axial regionalisation. This points to an asymmetry in developmental evolvability: early processes such as axial patterning via *hox* genes may be relatively labile due to redundancy and modularity, whereas later processes shaping vertebral morphology are more constrained, likely because they involve integrated growth and ossification.

Vertebral development in teleosts, including cichlids, begins with paired chondral condensations of the neural and haemal arches that form in an anterior-to-posterior sequence. The centra are subsequently ossified directly via intra-membranous ossification of the notochord sheath (Apschner et al., 2011; Woltering et al., 2018; Dietrich et al., 2020). The strong integration observed between precaudal and caudal vertebral shape (r-PLS = 0.886, Figure 3D) may therefore reflect this shared centrum ossification programme, whereby developmental changes affect all vertebrae derived from this process. In contrast, integration between precaudal vertebrae and pleural ribs is lower (r-PLS = 0.585, Figure 3A). In the haplochromine *Astatotilapia burtoni*, pleural ribs ossify from cartilaginous intermediates that form after centrum ossification (Woltering et al., 2018). This temporal and mechanistic distinction in developmental pathways may underlie the reduced integration between these skeletal elements. It nonetheless remains unclear when interspecific differences in vertebral and pleural rib shape arise during development, or indeed whether they emerge post-embryonically. As direct developers, cichlids hatch with a largely ossified vertebral column that may grow approximately isometrically with age (Fujimura and Okada, 2008) but patterns of skeletal and body scaling throughout ontogeny in cichlids remain poorly understood.

Mechanical forces arising from embryonic movement, musculoskeletal loading, and environmental factors such as swimming activity may contribute to shaping the vertebral column, raising the possibility that variation in activity throughout development could influence skeletal morphology. Evidence from laboratory, aquaculture, and wild fish consistently show that vertebral morphology is sensitive to mechanical loading. In zebrafish, vertebral shape changes as individuals mature from juveniles into adults, with repeated exposure to sustained water movement increasing centrum and bone volume as well as vertebral length (Nguyen et al., 2022; Suniaga et al., 2018). Similarly, studies in both cultured and wild fish indicate that mechanical stress associated with continued swimming can induce vertebral pathologies, including fractures, callus formation, and spinal deformities (Fjelldal et al., 2021). These effects appear to be especially pronounced during early ontogeny, where high levels of swimming activity and mechanical stress are linked to an increased incidence of spinal deformities (Bardon et al., 2009; Franz et al., 2025). While pathological, they nevertheless demonstrate that vertebral column development is responsive to mechanical loading in early and late ontogeny, raising the possibility that subtle variation in mechanical loading could contribute to non-pathological differences in vertebral morphology among species. A comparison of vertebral column development between divergent species would help determine when these differences first appear and whether they reflect early ossification differences or later remodelling.

Proportional increases in centrum size (Figure 4A, B), but not centrum elongation, are associated with whole-body elongation (see Figure 5). Proportional increases in centrum size may be necessary to maintain structural integrity, flexibility and effective force transmission along an elongate body axis. Body elongation in African cichlids is linked to predation, particularly active piscivory as in *Rhamphochromis* (Stiassny, 1981; Bucklow et al., 2025b), *but the underlying mechanics likely differ from those of other elongate predators such as barracuda (Sphyraena*) or Scombriformes, in which elongation is achieved through fewer, more elongate vertebrae that stiffen the body axis and facilitate sustained high-speed swimming (Mehta et al., 2010; Jimenez et al., 2023). In contrast, *Rhamphochromis* may retain greater axial flexibility while maintaining structural support, consistent with burst swimming or short-distance strike behaviours. However, this relationship is not universal: the rock-dwelling mbuna *Chindongo elongatus* exhibits the highest centrum aspect ratios despite having average vertebral counts (32 (Bucklow et al., 2026a)), and several mbuna species show similarly elevated ratios (Figure 5, black box). If increased centrum elongation corresponds to a stiffer axial column, this pattern is difficult to reconcile with the demands of navigating complex rocky habitats, where greater flexibility might be expected. Given the diversity within the radiation, integrating these morphological data with biomechanical measurements of swimming performance could clarify how axial skeleton structure, body shape, and locomotor function are linked (Satterfield et al., 2023).

In summary, our results highlight whole-body elongation as a key axis of morphological diversification in Lake Malawi cichlids, with changes in body elongation likely driving coordinated modifications across the axial skeleton rather than simple changes to individual vertebrae. This decoupling between vertebral count evolution and tightly integrated shape variation underscores the importance of considering multivariate morphological integration when interpreting the evolution of morphological traits. More broadly, these findings suggest that adaptive diversification in this radiation is facilitated by flexible developmental and functional systems capable of generating coordinated phenotypic change without requiring corresponding shifts in early developmental processes. Future work integrating comparative developmental data, functional biomechanics, and ecological context will be essential to determine how these integrated shape changes translate into performance differences and ecological specialisation, and whether similar principles underlie axial diversification in other vertebrate radiations.

## AUTHOR CONTRIBUTIONS

C.V.B, B.V and R.B. conceived the study. H.U. and C.V.B segmented vertebral models from *µ*CT-scans. K. C. provided the initial landmarking scheme (i.e. core 15 landmarks), C.V.B defined the additional landmarks to capture pleural rib shape and haemal spine positioning. H.U. landmarked all models. C.V.B performed the data analysis. C.V.B wrote the manuscript. B.V. and R.B. edited the manuscript. All authors reviewed the manuscript.

## ACKNOWLEDGMENTS

This research was supported by a Biotechnology and Biological Sciences Research Council (BBSRC) studentship (Grant No. 2445747), the John Fell Fund at the University of Oxford (Grant No. 0009780) and a European Research Council Starting Grant (No. 101163722, Counts). We thank Richard Durbin for the *µ*CT-scans of the specimens in the CAMZM collection used in this study that were *µ*CT-scanned at the Cambridge Biotomography Centre. The specimens were collected ethically under prescribed permits, and the results and data are published under an Access and Benefit Sharing agreement with the Government of Malawi. We acknowledge the contributions of Hannes Svardal, Karl Svardal, Richard Zatha, Bosco Rusuwa, George Turner and colleagues, and the Malawi Department of Fisheries and the Government of Malawi for their assistance in the collection of samples. We thank Liz Martin-Silverstone at the XTM Facility, Palaeobiology Research Group, University of Bristol for scanning the *Maylandia zebra* used in this study. Thank you to the fish whose lives were sacrificed for this work.

## SUPPLEMENTARY FIGURES

**Supplementary Figure 1.**
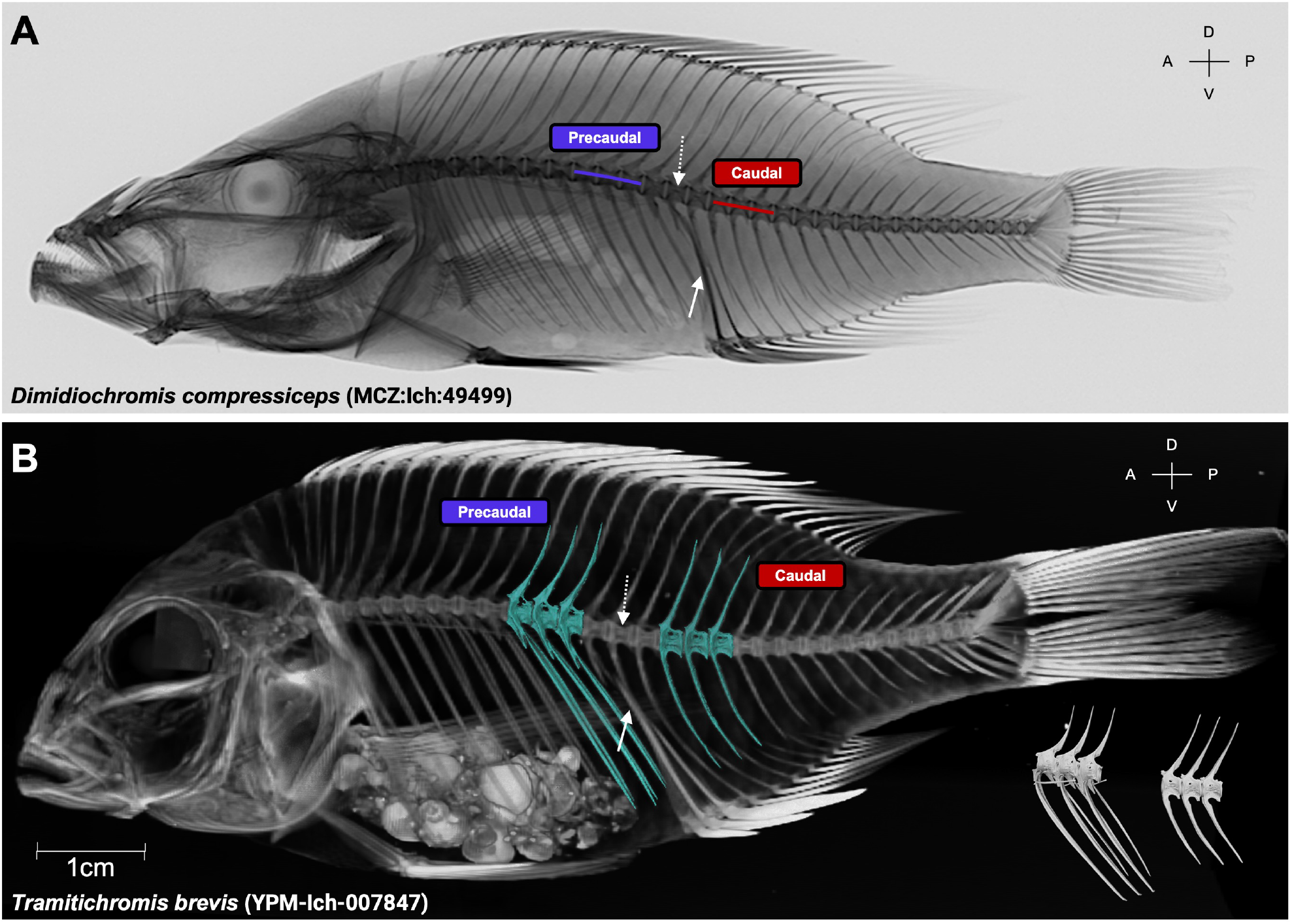
Segmentation of precaudal and caudal vertebrae. (A) Radiograph of *Dimidiochromis compressiceps* and (B) lateral reconstruction of a *µ*CT-scan of *Tramitichromis brevis*. Museum specimen numbers are indicated. Three precaudal (purple) and caudal (red) vertebrae were segmented from species according to their relative position to the first vertebrae (dashed white arrow) that articulates with the second anal fine spine (solid white arrow). The vertebrae immediately anterior and posterior to this vertebrae were ignored and the next three were segmented as precaudal and caudal vertebrae, respectively. For *Tramitichromis brevis*, the 3D models have been overlaid against the image reconstruction. Anatomical orientation axes are indicated (anterior–posterior, A–P; dorsal–ventral, D–V). Scale bar (1cm) is shown for *Tramitichromis brevis*.

**Supplementary Figure 2.**
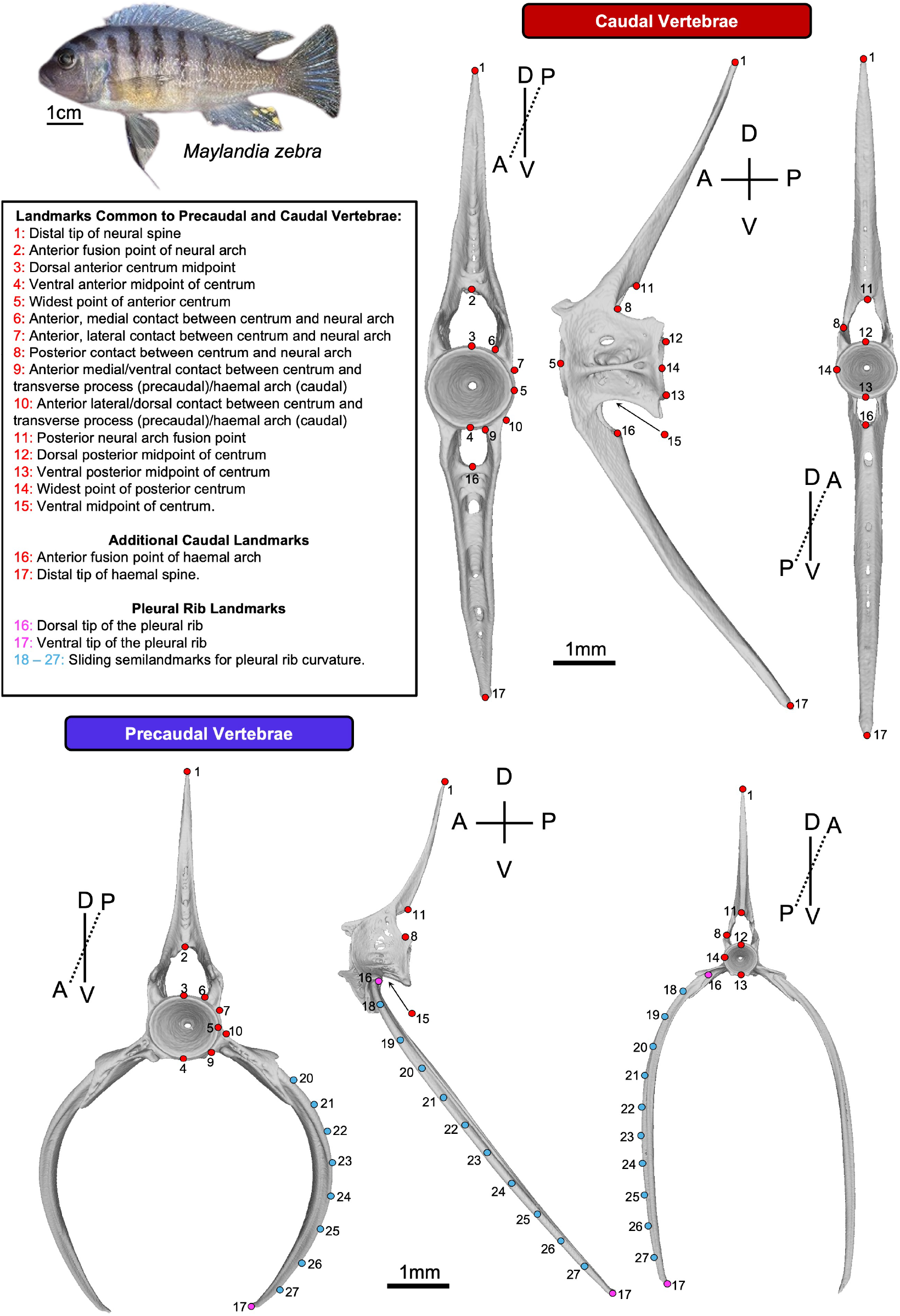
Landmark scheme. Example caudal (top) and precaudal (bottom) vertebrae from *Maylandia zebra* (top left) is indicated. Scale bar for *M. zebra* is 1cm. Red dots represent fixed, homologous landmarks. Landmarks 1-15 are homologous and shared between both precaudal and caudal vertebrae. Blue dots represent semi-landmarks of the pleural ribs that were permitted to slide, between two fixed landmarks added to the pleural ribs (pink). Precaudal and pleural ribs were landmarked together on the same 3D model despite being two independent bones. Two additional landmarks were added to the caudal vertebrae to capture shape variation in haemal spine shape. Anatomical orientation axes are indicated (anterior–posterior, A–P; dorsal–ventral, D–V). 1mm scale bars are indicated for the vertebrae.

